# Structure of RNA-dependent RNA polymerase from 2019-nCoV, a major antiviral drug target

**DOI:** 10.1101/2020.03.16.993386

**Authors:** Yan Gao, Liming Yan, Yucen Huang, Fengjiang Liu, Yao Zhao, Lin Cao, Tao Wang, Qianqian Sun, Zhenhua Ming, Lianqi Zhang, Ji Ge, Litao Zheng, Ying Zhang, Haofeng Wang, Yan Zhu, Chen Zhu, Tianyu Hu, Tian Hua, Bing Zhang, Xiuna Yang, Jun Li, Haitao Yang, Zhijie Liu, Wenqing Xu, Luke W. Guddat, Quan Wang, Zhiyong Lou, Zihe Rao

## Abstract

A novel coronavirus (2019-nCoV) outbreak has caused a global pandemic resulting in tens of thousands of infections and thousands of deaths worldwide. The RNA-dependent RNA polymerase (RdRp, also named nsp12), which catalyzes the synthesis of viral RNA, is a key component of coronaviral replication/transcription machinery and appears to be a primary target for the antiviral drug, remdesivir. Here we report the cryo-EM structure of 2019-nCoV full-length nsp12 in complex with cofactors nsp7 and nsp8 at a resolution of 2.9-Å. Additional to the conserved architecture of the polymerase core of the viral polymerase family and a nidovirus RdRp-associated nucleotidyltransferase (NiRAN) domain featured in coronaviral RdRp, nsp12 possesses a newly identified β-hairpin domain at its N-terminal. Key residues for viral replication and transcription are observed. A comparative analysis to show how remdesivir binds to this polymerase is also provided. This structure provides insight into the central component of coronaviral replication/transcription machinery and sheds light on the design of new antiviral therapeutics targeting viral RdRp.

**One Sentence Summary:** Structure of 2019-nCov RNA polymerase.

---

Corona Virus Disease 2019 (COVID-19) caused by a novel coronavirus 2019-nCoV has recently emerged (*1–3*). According to the World Health Organization (WHO) on March 11, 2020, 11800 infections and 4291 fatalities have been confirmed globally. 2019-nCoV is reported to be a new member of the betacoronavirus genus and is closely related to severe acute respiratory syndrome coronavirus (SARS-CoV) and to several bat coronaviruses (*4*). Compared to SARS-CoV and MERS-CoV, 2019-nCoV appears to exhibit faster human-to-human transmission, thus leading to the WHO declaration of a Public Health Emergency of International Concern (PHEIC)(*1, 3*).

CoVs employ a multi-subunit replication/transcription machinery, being assembled by a set of non-structural proteins (nsp) produced as cleavage products of the ORF1a and ORF1ab viral polyproteins (*5*) to facilitate virus replication and transcription. A key component, the RNA-dependent RNA polymerase (nsp12), catalyzes the synthesis of viral RNA and thus plays a central role in the replication and transcription cycle of 2019-nCoV, possibly with the assistance of nsp7 and nsp8 as co-factors (*5, 6*). Therefore, nsp12 is a primary target for the nucleotide analog antiviral inhibitors, *e.g.* remdesivir which shows potential to treat 2019-nCoV infections (*7, 8*). To inform drug design we have determined the structure of nsp12, in complex with its cofactors nsp7 and nsp8 by cryo-Electron Microscopy (Cryo-EM) using two different protocols, one in the absence of DTT (Dataset-1) and the other in the presence of DTT (Dataset-2).

The bacterially expressed full-length 2019-nCoV nsp12 (residues S1-Q932) was incubated with nsp7 (residues S1-Q83) and nsp8 (residues A1-Q198), and the complex was then purified (fig. S1). Cryo-EM grids were prepared using this complex and preliminary screening revealed excellent particle density with good dispersity. After collecting and processing 7,994 micrograph movies, we obtained a 2.9-Å resolution 3D reconstruction of one nsp12 monomer in complex with one nsp7-nsp8 pair and one nsp8. Like that observed in SARS-CoV nsp12-nsp7-nsp8 (*6*), we observed classes corresponding to an nsp12 monomer in complex one nsp8, as well as individual nsp12s, but these do not give atomic resolution reconstructions (fig. S2). However, the nsp12-nsp7-nsp8 complex provides all the structural information for complete and comprehensive analysis.

The structure of the 2019-nCoV nsp12 contains a “right hand” polymerase domain (residues S367-F920), and a nidovirus-unique N-terminal extension domain (residues D60-R249) adopting a nidovirus RdRp-associated nucleotidyltransferase (NiRAN)(*9*) architecture, being connected by an interface domain (residues A250-F369) (Figs. 1, A and B). An additional N-terminal β-hairpin (V31-K50) was built with the guidance of an unambiguous cryo-EM map (fig. S3A), inserting into the groove clamped by the NiRAN domain and the palm subdomain in the RdRp domain (Fig. 2). The nsp7-nsp8 pair shows a conserved structure similar to the SARS-CoV nsp7-nsp8 pair (*6, 10*). The orientation of the N-terminal helix of another individual nsp8 bound to nsp12 is shifted compared with that in the nsp7-nsp8 pair (fig. S4). The polypeptides of nsp12 can be traced in the map, except for S1-V30, T51-S68, H75, K103-V111, D336, T896-S904, F921-Q932, which cannot be observed.

**Fig. 1.**
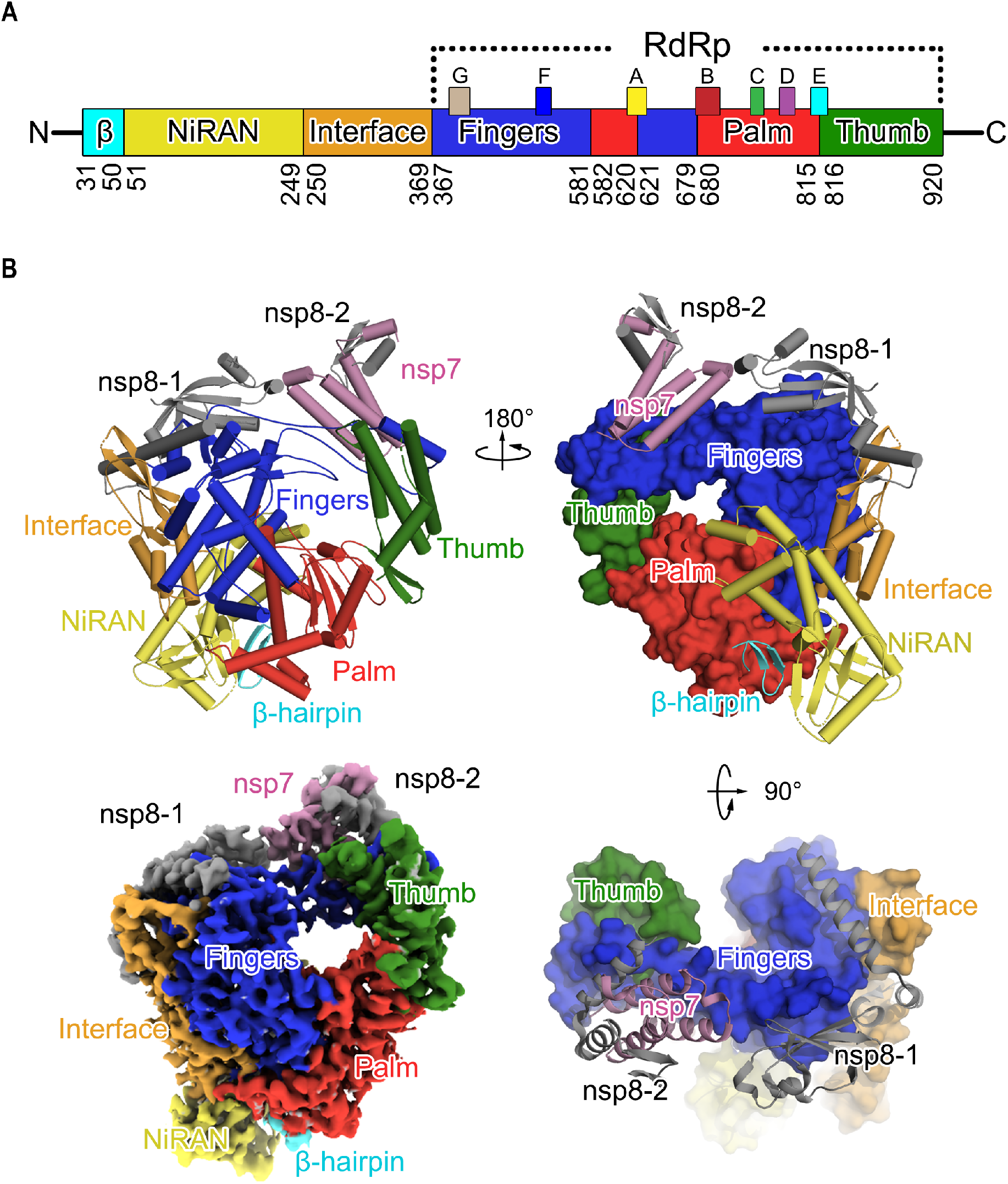
Structure of 2019-nCoV nsp12-nsp7-nsp8 complex. **(A)** Domain organization of 2019-nCoV nsp12. The interdomain borders are labeled with residue numbers. The N-terminal portion with no cryo-EM map density and the C-terminal residues that cannot be observed in the map are not included in the assignment. The polymerase motifs (A-G) have the same color scheme used in Fig. 3g. **(B)** Ribbon diagram of 2019-nCoV nsp12 polypeptide chain in three perpendicular views. Domains are colored the same as in (**A**). The individual nsp8 (nsp8-1) bound to nsp12 and that in the nsp7-nsp8 pair (nsp8-2) are in grey; the nsp7 is in pink. The bottom left panel shows an overview of the cryo-EM reconstruction of the nsp12-nsp7-nsp8 complex.

**Fig. 2.**
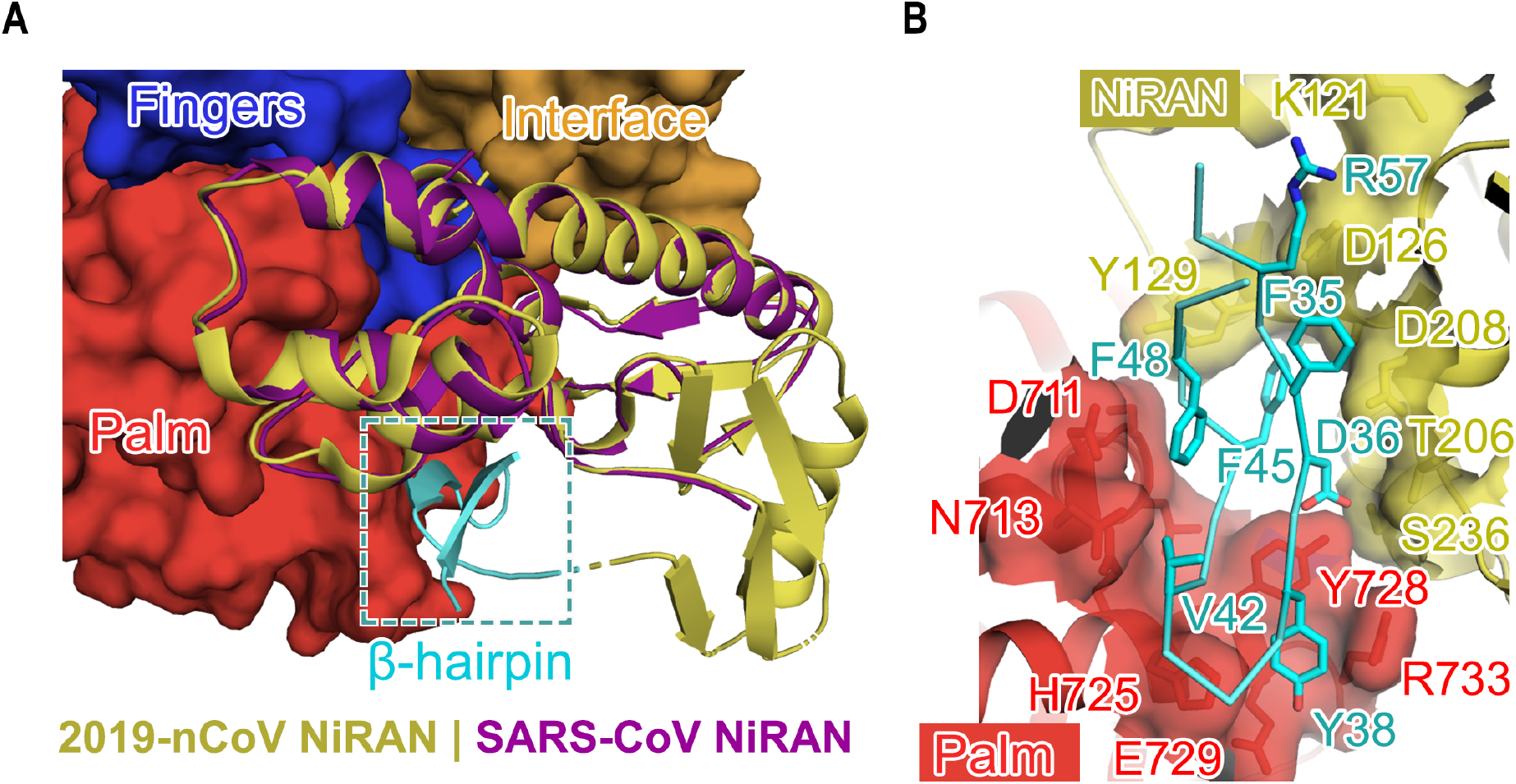
Structure of N-terminal NiRAN domain and β-hairpin. **(A)** Overall structure of the N-terminal NiRAN domain and β-hairpin of 2019-nCoV nsp12. The N-terminal NiRAN domain and β-hairpin of 2019-nCoV nsp12 are shown as yellow and cyan cartoons, while the other regions of 2019-nCoV nsp12 are shown as a molecular surface with the same color scheme used in Fig. 1. The NiRAN domain of SARS-CoV nsp12 is superimposed to its counterpart in 2019-nCoV nsp12 and is shown in purple. **(B)** Key interactions between β-hairpin and other domains. The β-hairpin is shown as a cyan tube with its key residues in stick mode. These have the closest contacts with other domains of 2019-nCoV nsp12. The interacting residues in the palm and fingers subdomain of the RdRp domain, and the NiRAN domain, are identified by the labels.

Although the overall architecture of 2019-nCoV nsp12-nsp7-nsp8 complex is generally similar to that of SARS-CoV with an r.m.s.d value of 0.82 for 1,078 Cɑ atoms there are key features that distinguish them. First, the cryo-EM map allows us to build the structure of 2019-nCoV nsp12 covering almost all residues, presenting the first insight into the complete architecture of coronaviral RdRp. In SARS-CoV nsp12, there are seven helices with a three-stranded β-sheet at the end that make up the NiRAN domain (*6*) (Fig. 2A). Residues Y69-R118 constitute an additional structural block with three anti-parallel β-strands and one helix. Residues N215-D218 form a β-strand in 2019-nCoV nsp12. We reasoned that the contact of this region with the strand (residues V96-A100) helps to stabilize it. As a result, these four strands form a compact semi β-barrel architecture. Therefore, we identify residues Y69-R249 as the complete coronaviral NiRAN domain. Second, the cryo-EM map helps us to identify a unique N-terminal β-hairpin (Figs. 1A and 2A). This β-hairpin inserts in the groove clamped by the NiRAN domain and the palm subdomain in the RdRp domain and forms a set of close contacts to stabilize the overall structure (Fig. 2B and fig. S5). Another point to note is that, we have observed C301-C306 and C487-C645 form disulfide bonds in the absence of DTT (Dataset-1). However, in the presence of DTT (Dataset-2), chelated zinc ions are present and in the same location as that observe in SARS-CoV (fig. S3B). We reasoned the different purification buffer has led to this variation.

The polymerase domain adopts the conserved architecture of the viral polymerase family (*11*) and is composed of three subdomains, including a fingers subdomain (residues S397-A581 and K621-G679), a palm subdomain (residues T582-P620 and T680-Q815), and a thumb subdomain (residues H816-E920) (Fig. 1). The catalytic cations, which are usually observed in viral polymerase structure for RNA synthesis (*12, 13*) are not observed, possibly because manganese was not included in the buffer for purification or in the cryo-EM sampling buffer.

The active site chamber of 2019-nCoV RdRp domain is formed by the conserved polymerase motifs A-G in the palm domain and configured like other RNA polymerases (Figs. 1A and 3A; fig. S6). Motif A covers residues 611-TPHLMGW**D**YPKCDRAM-626 with a potential divalent-cation-binding residue D618, which is conserved in HCV ns5b (residue D220) and poliovirus (PV) 3D^pol^ (residue D233) (*12, 13*) (Figs. 3, B and C). Motif B is in the region 680-TSSGDATTAYANSVFNICQAVTANVNALLST-710. Motif C (residues 753-FSMMIL**SDD**AVVCFN-767) contains the catalytic residues (759-**SDD**-761) in the turn between the two β-strands. These catalytic residues are also conserved in viral RdRp, e.g. 317-**GDD**-319 in HCV ns5b and 327-**GDD**-329 PV 3D^pol^ with the first residue being a glycine residue as an alternative. Motif D is assigned as 775-LVASIKNFKSVLYYQNNVFMSE-796. Motif E (810-HEFCSQHTMLV-820) forms the tight loop. Motif F (912-KKNQHGGLRE-921) forms an ordered loop (β-loop) connecting two anti-parallel β-strands. Motif G is in 500-KSAGFPFNKWGKARLYYDS-518.

**Fig. 3.**
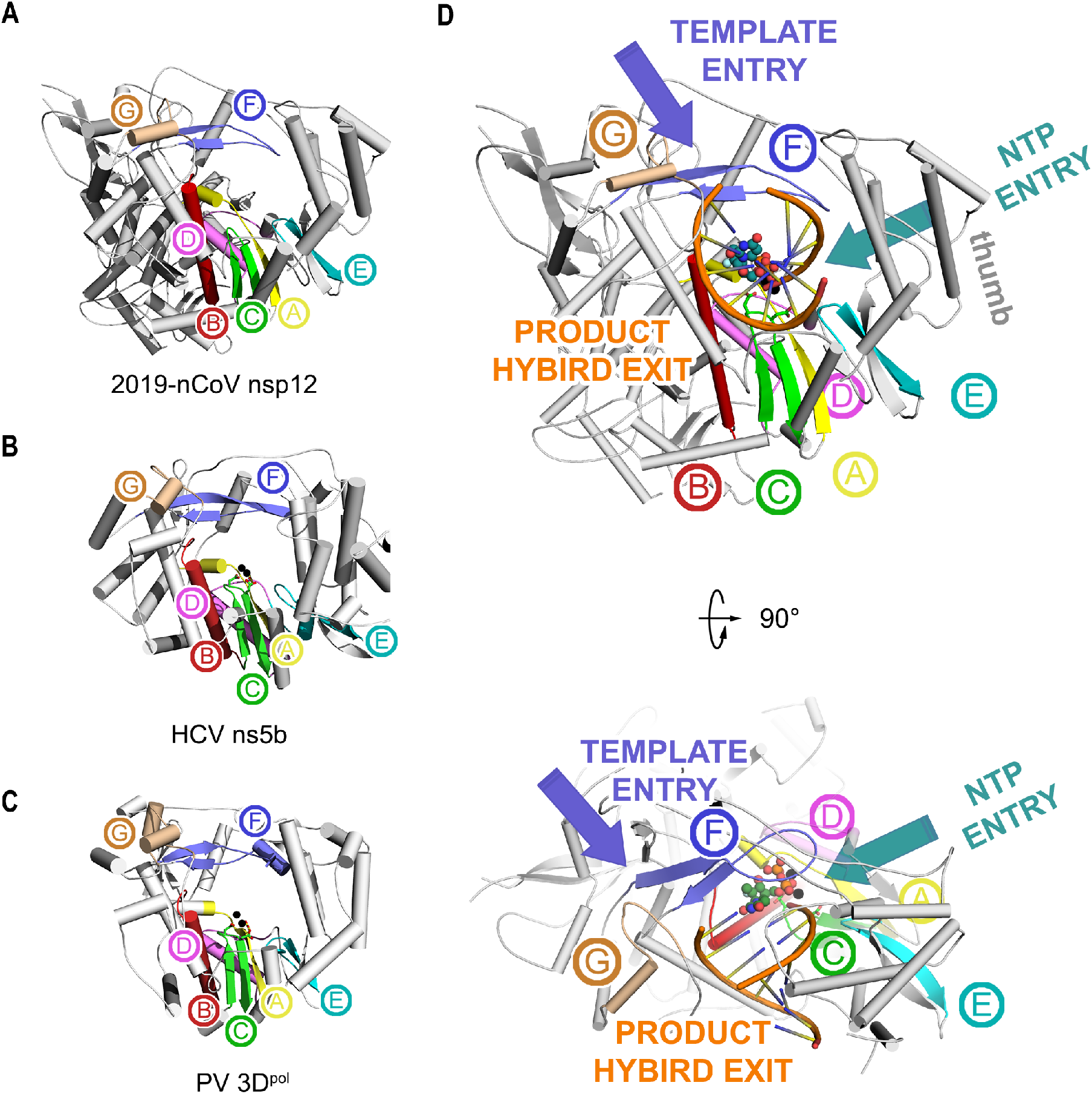
The RdRp core region. **(A-C)** Structural comparison of 2019-nCoV nsp12 **(A)**, HCV ns5b (PDB ID: 4WTG)(*12*) **(B)**, and PV 3D^pol^ (PDB ID: 3OLB)(*13*) **(C)**. The three structures are displayed in the same orientation. The polymerase motifs are colored as: motif A, yellow; motif B, red; motif C, green; motif D, violet; motif E, cyan; motif F, blue; and motif G, light brown. **(D)** The template entry, NTP entry, product hybrid exit paths in 2019-nCoV nsp12 are labeled with slate, deep teal and orange colors. Two catalytic manganese ions (black spheres), pp-sofosbuvir (dark green spheres for carbon atoms) and primer template (orange) from the structure of HCV ns5b in complex pp-sofosbuvir (PDB ID: 4WTG)(*12*) are superposed to 2019-nCoV nsp12 to indicate the catalytic site and nucleotide binding position.

In this structure, the four positively charged solvent-accessible paths converge into a central cavity where the RdRp motifs mediate template-directed RNA synthesis (Fig. 3D). The configurations of the template/primer entry paths, the nucleoside triphosphate (NTP) entry channel, and the nascent strand exit path are similar to those described for HCV and PV polymerase (*14*) (Figs. 3, B and C). The NTP entry channel is formed by a set of hydrophilic residues, including K545, R553 and R555 in motif F. The RNA template is expected to enter the active site composed of motifs A and C through a groove clamped by motif F and G. Motif E and the thumb subdomain support the primer strand. The product-template hybrid exits the active site through the RNA exit tunnel at the front side of the polymerase.

Remdesivir, the single *Sp* isomer of the 2-ethylbutyl L-alaninate phosphoramidate prodrug (*15*) (fig. S7), has been reported to inhibit 2019-nCoV proliferation and have potential to treat patients in the clinic (*7, 8*). The efficacy of chain-terminating nucleotide analogs requires viral RdRps to recognize and successfully incorporate the active form of the inhibitors into the growing RNA strand. Because of the structural conservation on the polymerase catalytic chamber between 2019-nCoV and HCV ns5b polymerase, as well as the likely similar mechanisms of action of remdesivir and sofosbuvir (*12, 16*) (2’-F-2’-C-methyluridine monophosphate prodrug which targets HCV ns5b and has been approved for the treatment of chronic HCV infection) (fig. S7), a model of 2019-nCoV nsp12 with remdesivir diphosphate is proposed (Fig. 4). In 2019-nCoV nsp12, the 2’ hydroxyl of the incoming NTP is likely to form hydrogen bonds with T680 and N691 in motif B (Figs. 4, A and B). In addition, D623 in motif A is positioned to interrogate the 3’ hydroxyl through hydrogen bonding. Moreover, the hydrophobic side chain of V557 in motif F is likely to stack with and stabilize the +1 template RNA uridine base to base pair with the incoming triphosphate remdesivir (ppp-remdesivir). In the structures of HCV ns5b elongation complex or its complex with pp-sofosbuvir, a key feature is that the incorporated pp-sofosbuvir disrupts the hydrogen bonding network between S282 and D225 (Figs. 4, C and D), which is necessary to stabilize the incoming natural nucleotide (*12*). However, key contacts formed by S282 with the incoming nucleotide and the surrounding environment can result in an S→T resistance mutation that is already observed in the clinic (*17*). In the structure of apo 2019-nCoV nsp12, the orientation of S682 and D623, which are strictly conserved with S282 and D225 in HCV ns5b, is similar to that observed in the HCV ns5b-pp-sofosbuvir complex, indicating a different recognition mechanism with that inhibitor. Nevertheless, the close distance of S682 with the incoming ppp-remdesivir and the environment surrounding the inhibitor alters the appearance of a drug-resistant mutation associated with remdesivir treatments.

**Fig. 4.**
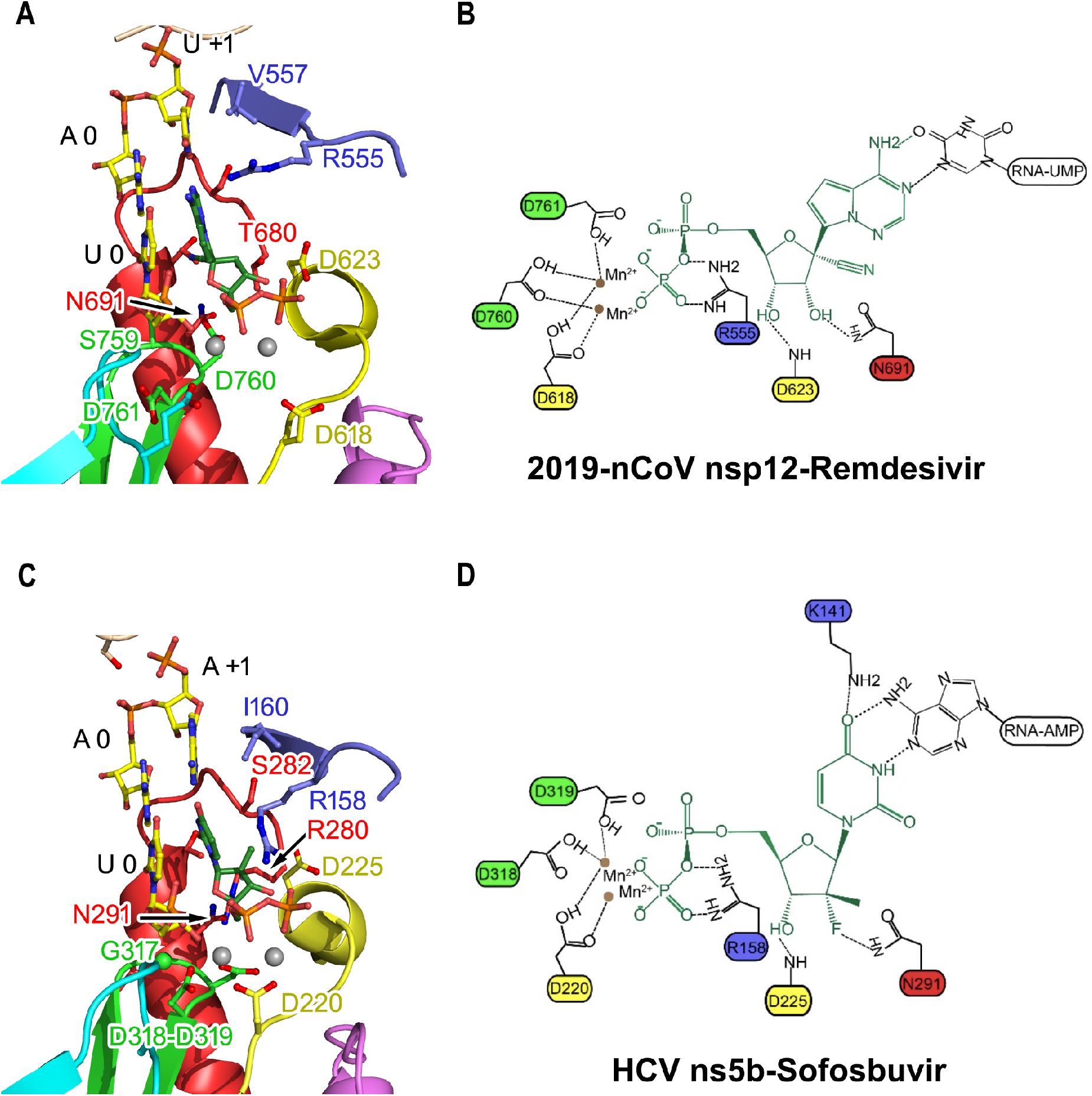
Incorporation model of remdesivir in 2019-nCoV nsp12. The polymerase motifs are colored as in Fig. 3. Superposition of the structure of HCV ns5b in complex with pp-sofosbuvir (PDB ID: 4WTG)(*12*) shows the positions of the two catalytic manganese ions (brown spheres), the priming nucleotide (U 0), template strand, and the incoming pp-remdesivir **(A, B)** or pp-sofosbuvir **(C, D)**.

The rapid global spread of 2019-nCoV has emphasized the need for the development of new coronavirus vaccines and therapeutics. The viral polymerase nsp12 looks an excellent target for new therapeutics, especially given that lead inhibitors already exist in the form of compounds such as remdesivir. This target, in addition to other promising drug targets such as the main protease, whose structure was recently determined by our group (*18*) in complex with an inhibitor (*19, 20*) will support the development of a cocktail of anti-coronavirus treatments that potentially can be used for the discovery of broad-spectrum antivirals.

## Acknowledgments

We would like to pay an exceptional tribute to ShanghaiTech University and their administrative team, as well as the Bio-Electron Microscopy Facility for their great care and support to our research team to enable us to carry out this research in a safe and healthy environment. It would have been impossible for us to attain this achievement without their tremendous efforts in the last two months during the nCoV-19 pandemic. We also would like to convey our special thanks to Tsinghua University for their exceptional permission to allow five of Z. R.'s students to go back to the laboratory to prepare the protein samples for this study. We also must express our gratitude to the campus service team of ShanghaiTech University scientific research platform of Shanghai Institute for Advanced Immunochemical Studies (SIAIS), and National Center for Protein Science Shanghai (NCPSS), as well as all the manager and technician individuals those who provided onsite or remote technical support. Their kind help and fearless support are pivotal to this work during the epidemic. We would like to thank the University of Queensland and Diamond Light Source for their collaboration.

## Author contributions

Z.R. conceived, initiated, and coordinated the project. Y.H, L.Y., Yao.Z., H.W., Yan.Z., C.Z., Z.M., L.Z., J.G., L.Z., Yi.Z. and Tianyu.H. purified the protein; L.Y. supervised sample purification; Y.G., F.L., L.C., Q.S. and Tian.H. collected the cryo-EM data; Y.G., F.L. and T.W. processed cryo-EM data; Q.W., L.Y., Y.G. built and refined the structure model; and the manuscript was written by Z.L., Q.W., L.Y., Y.G., F.L., L.G. and Z.R.. All authors discussed the experiments and results, read and approved the manuscript;

## Data and materials availability

The cryo-EM maps and the structures were deposited into the Electron Microscopy Data Bank (EMDB) and Protein Data Bank (PDB) with the accession numbers EMD-30127 and PDB 6M71 for Dataset-1, EMD-30178 and PDB 7BTF for Dataset-2 (under the reducing condition).

## Supplementary Materials

Materials and Methods

Figures S1-S7

Tables S1

References (*21–28*)

## Supplementary Materials

### Materials and Methods

#### Protein production and purification

The 2019-nCoV nsp12 (GenBank: MN908947) gene was cloned into a modified pET-22a vector, with the C-terminus possessing a 10 × His-tag. The plasmids were transformed into *E. coil* BL21 (DE3), and the transformed cells were cultured at 37 °C in LB media containing 100 mg/L ampicillin. After the OD_600_ reached 0.8, the culture was cooled to 16 °C and supplemented with 0.5 mM IPTG. After overnight induction, the cells were harvested through centrifugation, and the pellets were resuspended in lysis buffer (20 mM Tris-HCl, pH 8.0, 150 mM NaCl, 4 mM MgCl, 10% glycerol) and homogenized with an ultra-high-pressure cell disrupter at 4 °C. The insoluble material was removed through centrifugation at 12,000 rpm. The fusion protein was first purified by Ni-NTA affinity chromatography and elution with lysis buffer and then further purified by passage through a Hitrap Q ion-exchange column (GE Healthcare, USA) before loading onto a Superdex 200 10/300 Increase column (GE Healthcare, USA) in a buffer containing 20 mM Tris-HCl, pH 7.5, 250 mM NaCl and 4 mM MgCl_2_. Purified nsp12 was concentrated to 4 mg ml/L and stored at 4 °C.

The full-length 2019-nCoV nsp7 and nsp8 were co-expressed in *E. coil* BL21 (DE3) cells as a no-tagged protein and a 6 × His-SUMO fusion protein, respectively. After purification by Ni-NTA (Novagen) affinity chromatography, the nsp7-nsp8 complex was eluted through on-column tag cleavage by ULP protease. The eluate was further purified by Hitrap Q ion-exchange column (GE Healthcare, USA) and a Superdex 200 10/300 Increase column (GE Healthcare, USA) in a buffer containing 20 mM Tris-HCl, pH 7.5, 250 mM NaCl and 4 mM MgCl_2_.

#### Cryo-EM sample preparation and data collection

An aliquot of 3 μL of the sample at 0.7 mg/mL (added with 0.025% DDM) was applied onto a H_2_/O_2_ glow-discharged, 300-mesh Quantifoil R1.2/1.3 grid (Quantifoil, Micro Tools GmbH, Germany). The grid was then blotted for 3.0 s with a blot force of 0 at 8°C and 100% humidity and plunge-frozen in liquid ethane using a Vitrobot (Thermo Fisher Scientific, USA). Cryo-EM data were collected with a 300 keV Titan Krios electron microscope (Thermo Fisher Scientific, USA) and a K2 Summit direct electron detector (Gatan, USA). Images were recorded at EFTEM with a 165000× magnification and calibrated super-resolution pixel size 0.82 Å/pixel. The exposure time was set to 5 s with a total accumulated dose of 60 electrons per Å^2^. All images were automatically recorded using SerialEM (*21*). For Dataset-1, a total of 7,994 images were collected with a defocus range from 1.0 μm to 1.8 μm. For Dataset-2 (under reducing conditions), a total of 8494 images were collected with a defocus range from 1.1 μm to 2.0 μm. Statistics for data collection and refinement are in Table S1.

#### Cryo-EM image processing

All dose-fractioned images were motion-corrected and dose-weighted by MotionCorr2 software (*22*) and their contrast transfer functions were estimated by cryoSPARC patch CTF estimation. For Dataset-1, a total of 2,334,248 particles were auto-picked using blob picker and extracted with a box size of 300 pixels in cryoSPARC (*20*). The following 2D, 3D classifications and refinements were all performed in cryoSPARC. 918,133 particles were selected after two rounds of 2D classification. 100,000 particles were used to do Ab-Initio reconstruction in five classes, and then these five classes were used as 3D volume templates for heterogeneous refinement with all selected particles, with 110,176 particles converged into one class. Next, this particle set was used to perform homogeneous refinement, yielding a resolution of 3.1 Å. After local refinement, the final resolution reached 2.9 Å. For Dataset-2, the image processing was conducted using a similar pipeline. 753,481 particles were auto-picked initially, and 145,388 particles were selected after final heterogeneous refinement. The resolution reached 2.99 Å after non-uniform refinement and 2.95 Å after local refinement with a mask.

#### Model building and refinement

To solve the structure of the 2019-nCoV nsp12-nsp7-nsp8 complex, the structure of the SARS-CoV nsp12 (*24*) and nsp7-8 (*25*) were individually placed and rigid-body fitted into the cryo-EM map using UCSF Chimera (*26*). After the corresponding amino acids were replaced with those from 2019-nCoV, the model was manually built in Coot (*27*) with the guidance of the cryo-EM map, and in combination with real space refinement with Phenix (*28*). The data validation statistics are shown in Table S1.

---

**Fig. S1.**
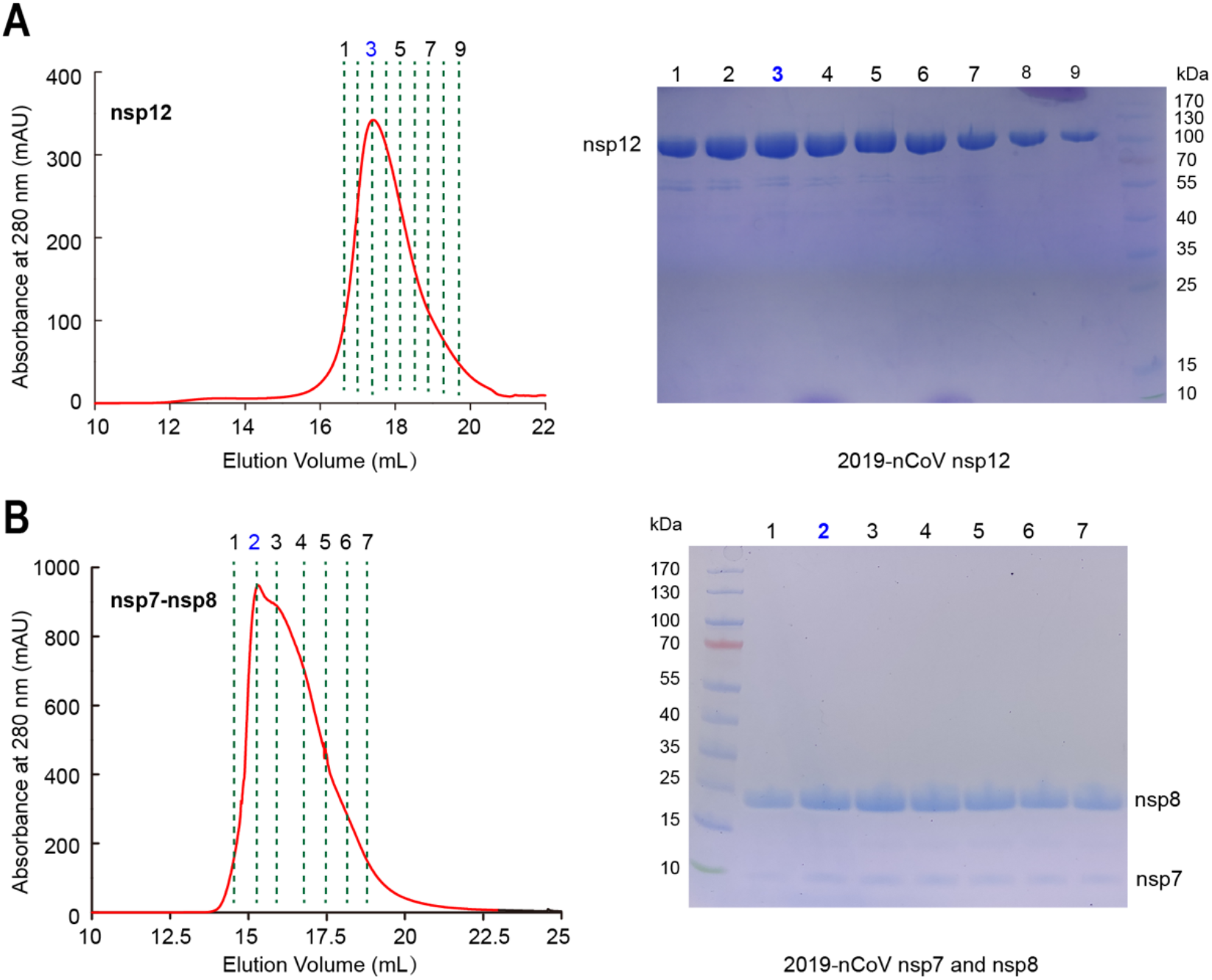
The purification of 2019-nCoV nsp12 and nsp12-nsp7-nsp8 complex. **(A)** Size-exclusion chromatogram of the affinity-purified the 2019-nCoV nsp12 and **(B)** nsp7-nsp8 complex. Data from the Superdex 200 10/30 column are shown in red. Seven fractions from the gel filtration peaks were pooled. The target proteins were analyzed by SDS-PAGE. Standard protein markers are shown in the unnumbered lane with molecular weights labelled.

**Fig. S2.**
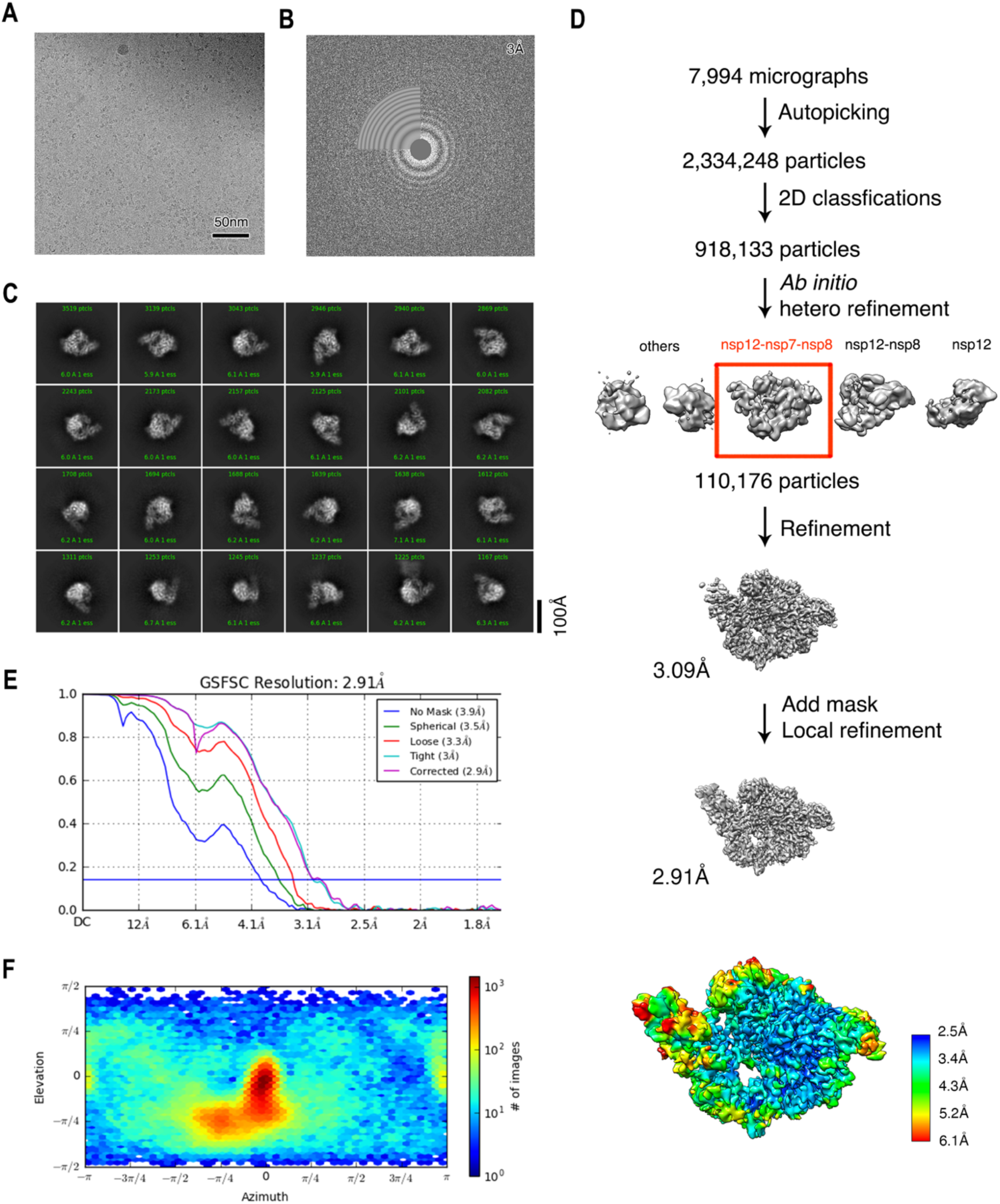
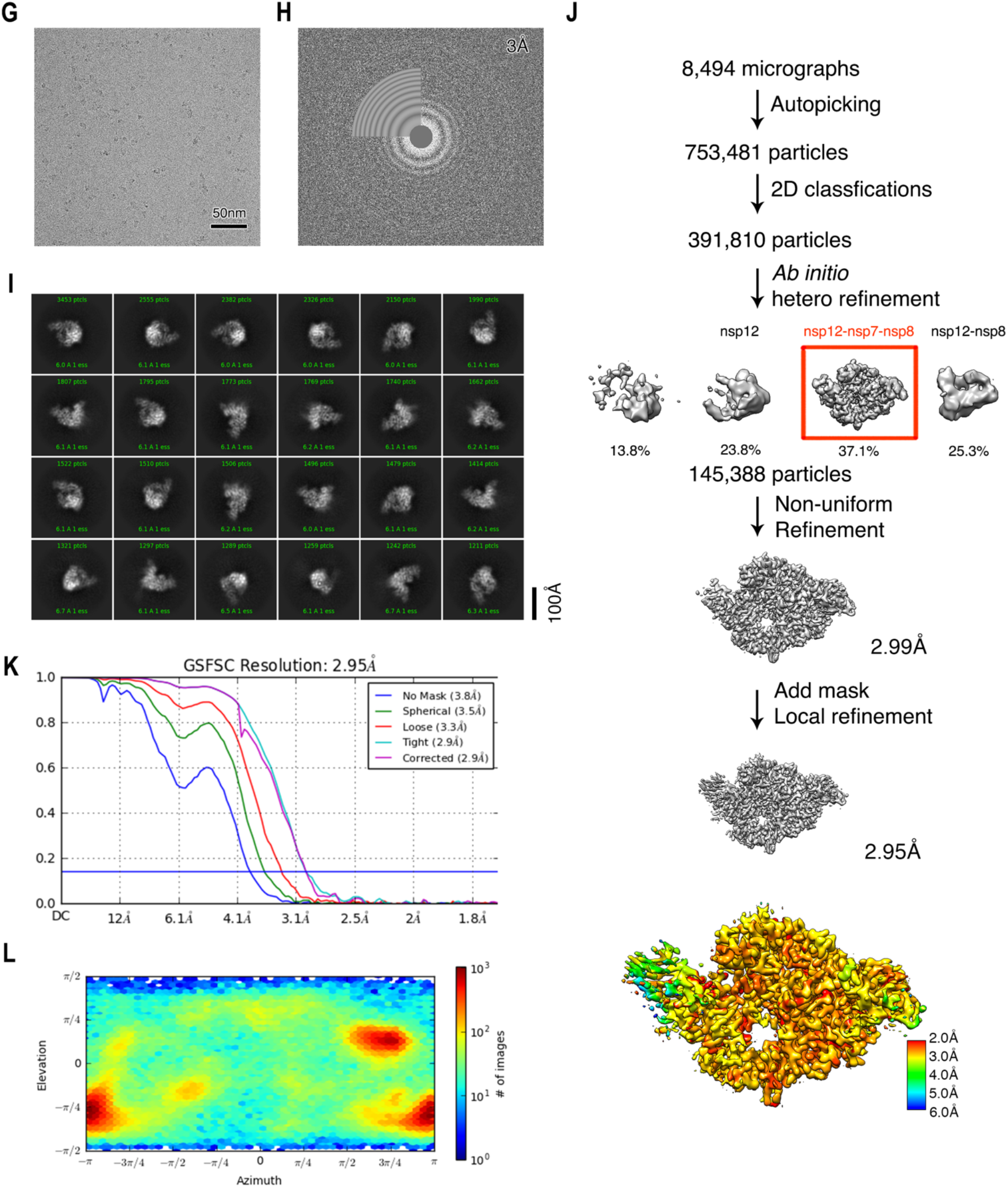
Cryo-EM reconstruction. **(A)** Raw image of the nsp12-nsp7-nsp8 complex particles in vitreous ice recorded at defocus values of −1.0 to −1.8 μm. Scale bar, 50 nm. **(B)** Power spectrum of the image shown in (A), with an indication of the spatial frequency corresponding to 3.0 Å resolution. **(C)** Representative class averages. The edge of each square is 246 Å. **(D)** The data processing scheme. Overview of nsp12-nsp7-nsp8 reconstruction is shown in the bottom panel with Local resolution. **(E)** Fourier shell correlation (FSC) of the final 3D reconstruction following gold standard refinement. FSC curves are plotted before and after masking. **(F)** Angular distribution heatmap of particles used for the refinement. **(G-I)** Data processing procedure and corresponding results for Dataset-2 (collected under reducing conditions).

**Fig. S3.**
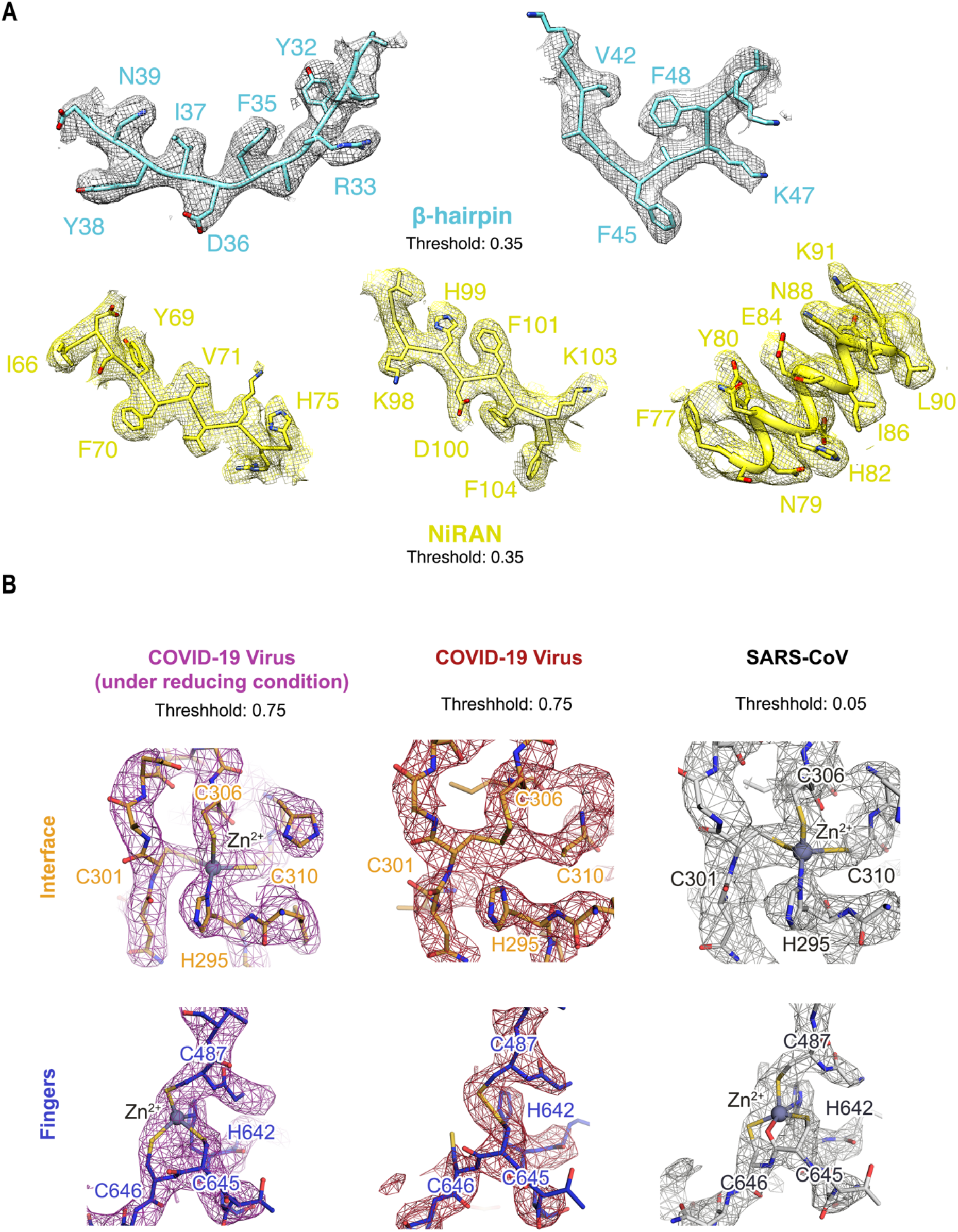
Cryo-EM map of β-hairpin (A) and disulfide bonds (B). **(A)** Structure and representative map of β-hairpin and NiRAN. **(B)** Raw cryo-EM map (mesh) for the nsp12-nsp7-nsp8 complex is shown in magenta and red (COVID-19 virus) for Dataset-2 and Dataset-1, respectively or grey (SARS-CoV, EMD-0520). Structures near the disulfide bond region of the Interface domain (orange) and Fingers domain (deep blue) are shown as stick models. The corresponding region in SARS-CoV (PDB ID: 6NUR) is shown on the right.

**Fig. S4.**
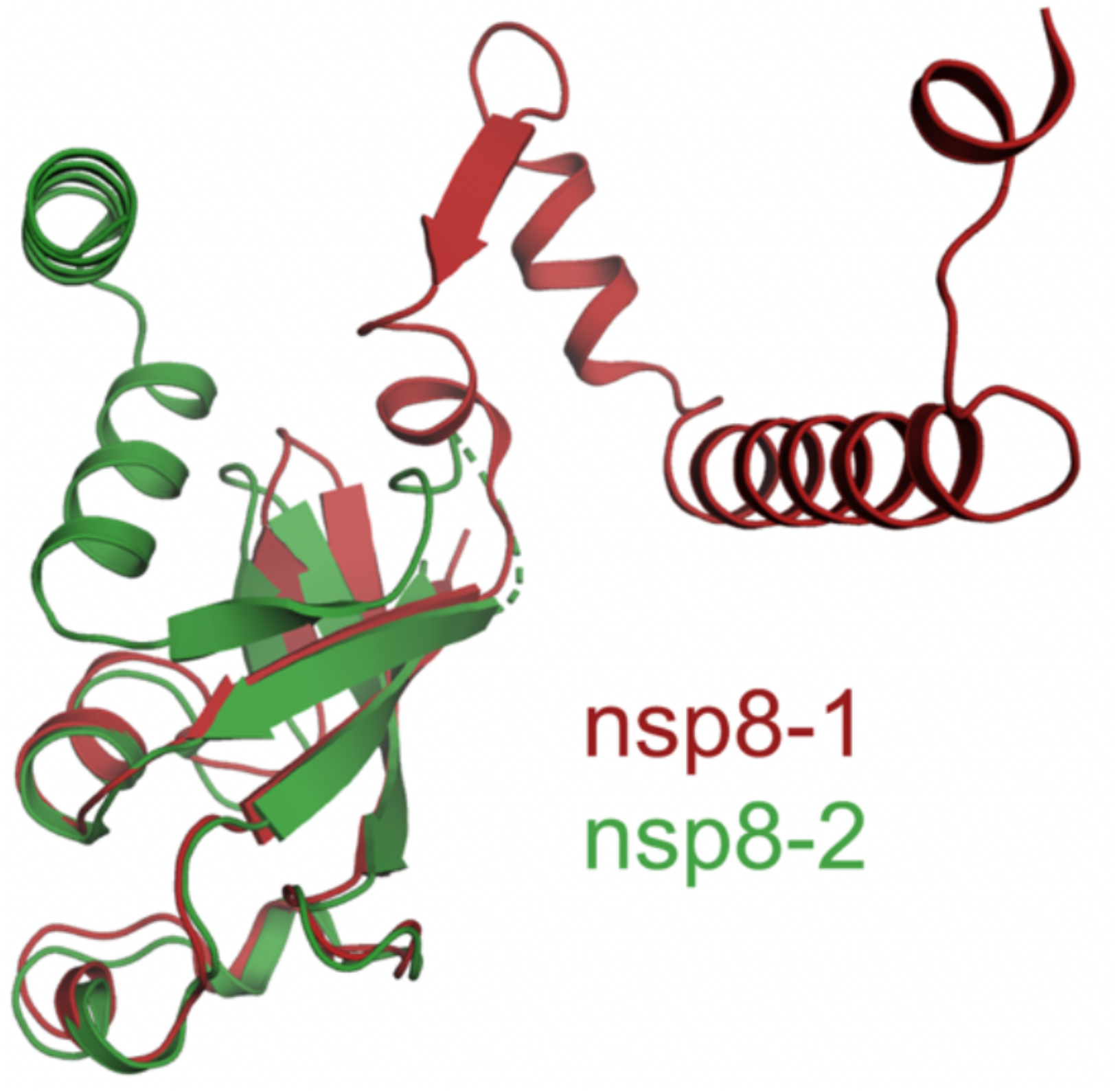
Structure comparison of two nsp8 molecules bound to 2019-nCoV nsp12. The nsp8-1 (in red) refers to the individual nsp8 molecule bound to nsp12. The nsp8-2 (in green) refers to the nsp8 molecule in nsp7-nsp8 pair. The two nsp8 molecules are aligned and shown in the same orientation.

**Fig. S5.**
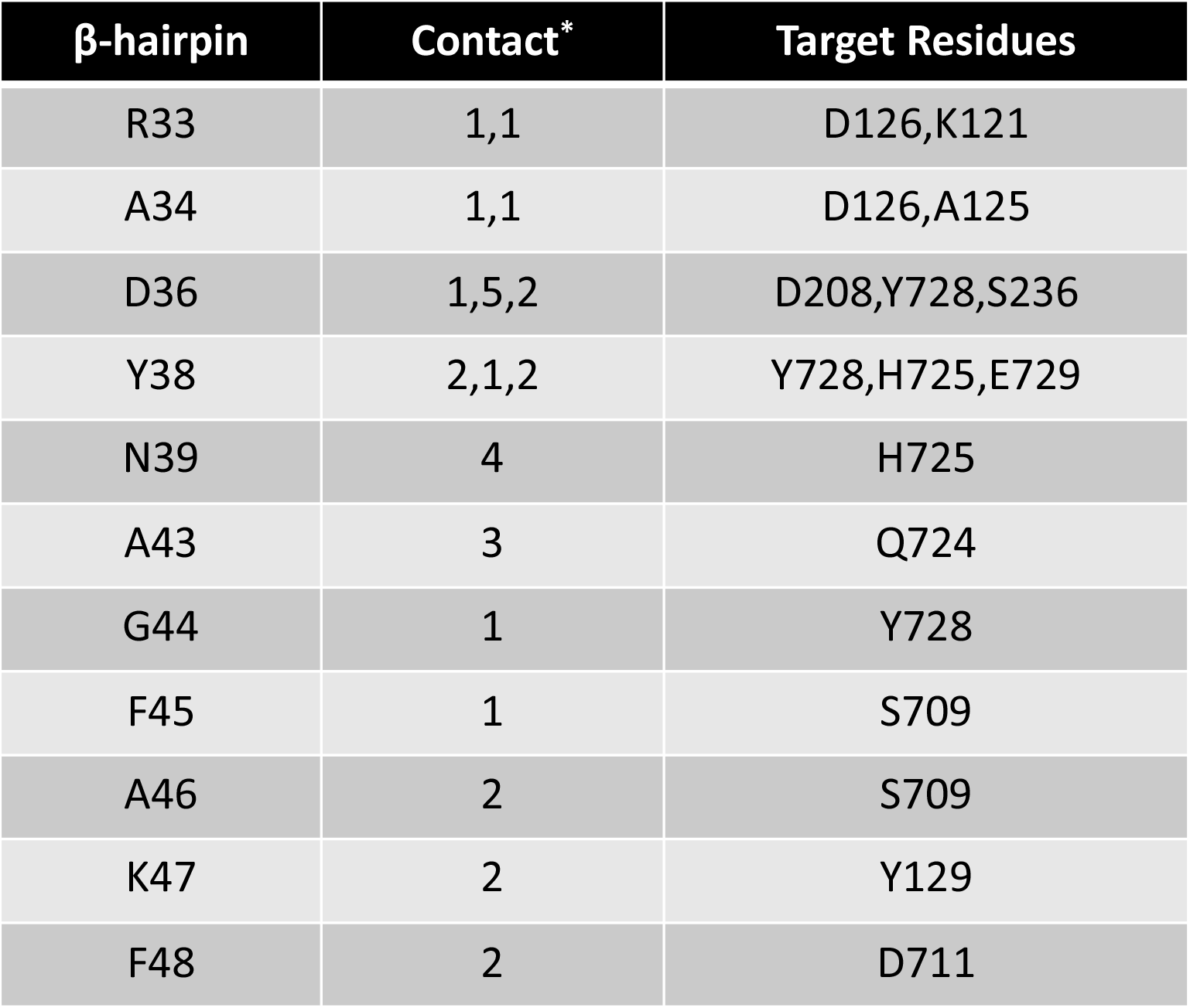
Interaction between the β-hairpin with other domains. *Numbers represent the number of atom-to-atom contacts between the residues of β-hairpin and the residues in other domains. These were analyzed by the Contact program in the CCP4 suite (with a distance cutoff of 3.5 Å).

**Fig. S6.**
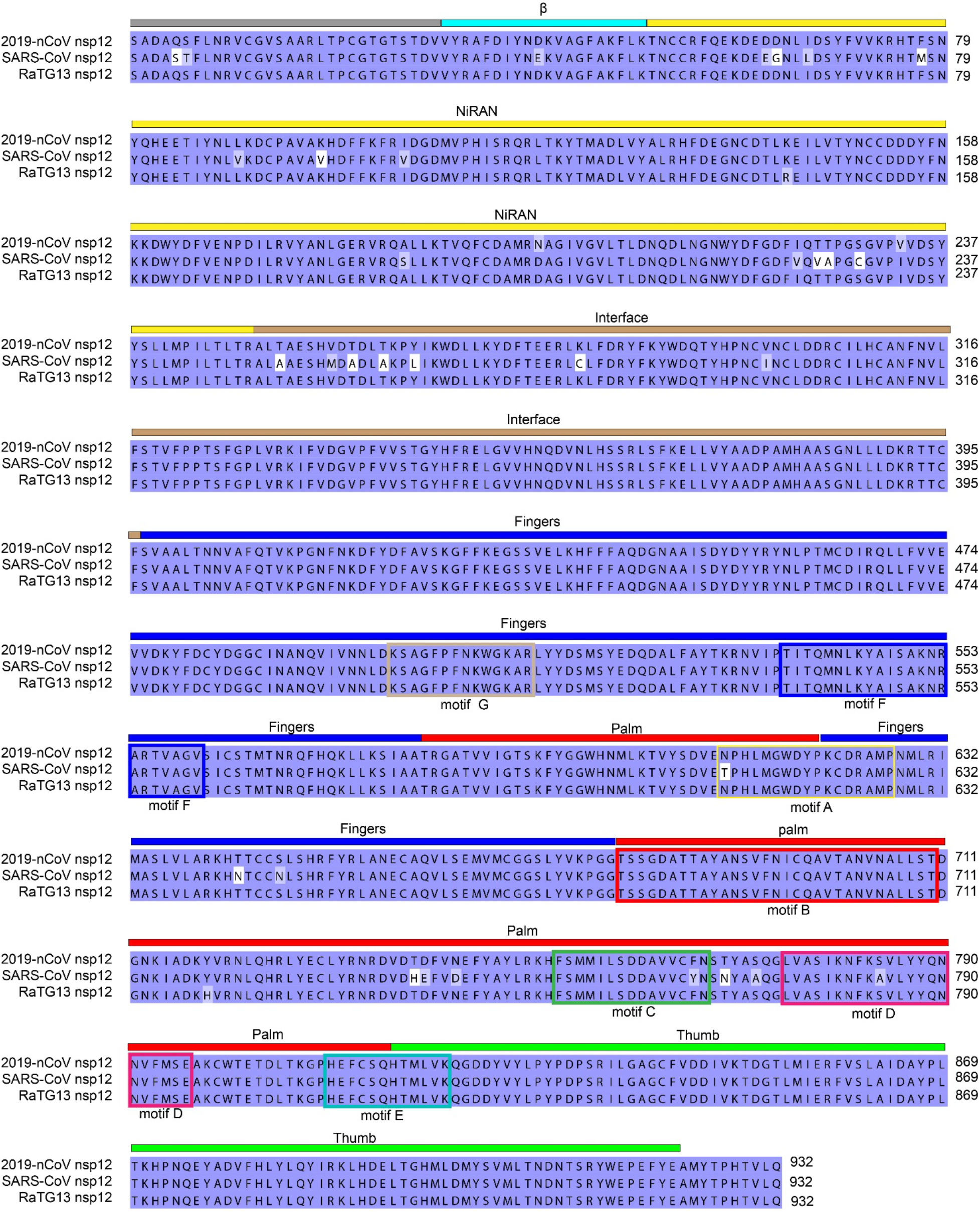
Sequence alignment of nsp12 proteins encoded by 2019-nCoV, SARS-CoV and RaTG13. The residues with blue, light blue or white backgrounds indicate the identical, conserved or non-conserved residues, respectively. Domain arrangement and key RdRp motifs are also indicated with the same color scheme as that in Fig. 1.

**Fig. S7.**
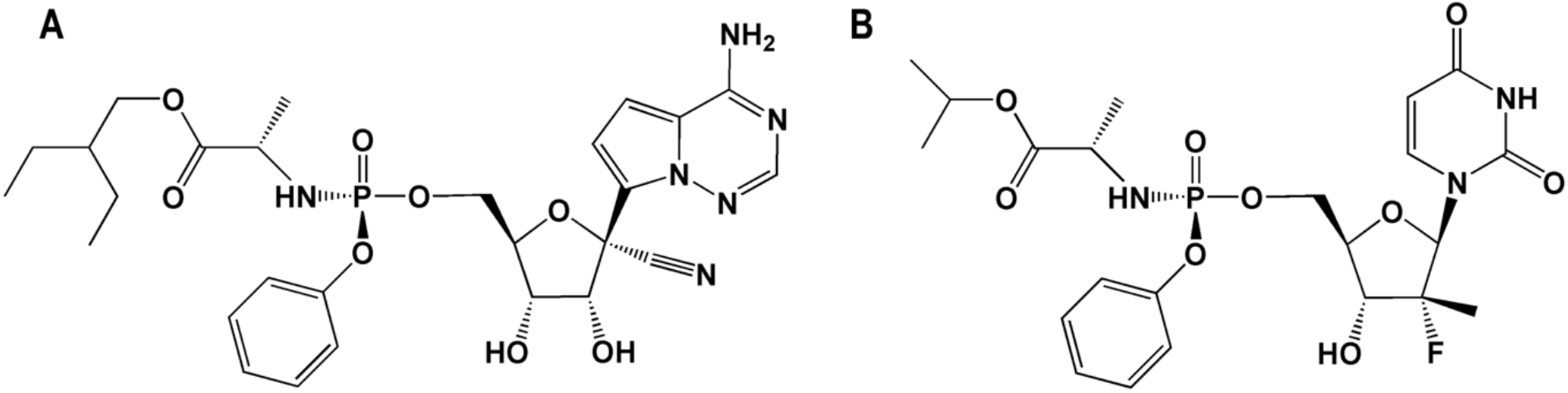
Chemical structures of (A) remdesivir and (B) sofosbuvir in their prodrug forms.

**Table S1.** Cryo-EM data statistics.

|  | COVID-19 Virus<br>nsp12-nsp7-nsp8<br>complex | COVID-19 Virus<br>nsp12-nsp7-nsp8 complex<br>under reducing condition |
| --- | --- | --- |
| **PDB entry** | 6M71 | 7BTF |
| **EMDB entry** | 30127 | 30178 |
| **Data collection and processing** |  |  |
| Magnification | 165,000 | 165,000 |
| Voltage (keV) | 300 | 300 |
| Electron exposure (e<sup>-</sup>/Å<sup>2</sup>) | 60.00 | 60.00 |
| Defocus range (μm) | -1.8 to -1.0 | -2.0 to -1.1 |
| Pixel size (Å) | 0.82 | 0.82 |
| Symmetry imposed | C1 | C1 |
| Initial particle images (no.) | 2,334,248 | 753,481 |
| Final particle images (no.) | 110,176 | 145,388 |
| Map global resolution (Å) | 2.91 | 2.95 |
| Global resolution FSC threshold | 0.143 | 0.143 |
| Map local resolution range (Å) | 1.8-7.8 | 1.8-7.5 |
| Local resolution FSC threshold | 0.143 | 0.143 |
| **Refinement** |  |  |
| Model resolution (Å) | 2.9 | 2.9 |
| FSC threshold | 0.143 | 0.143 |
| Model resolution range (Å) | ∞ to 2.9 | ∞ to 2.9 |
| Map sharpening *B* factor (Å<sup>2</sup>) | 80.2 | 97.6 |
| Model composition |  |  |
| Non-hydrogen atoms | 8,550 | 9,783 |
| Protein residues | 1,077 | 1,227 |
| Ligands | 0 | 2 |
| *B* factors (Å<sup>2</sup>) |  |  |
| Protein | 67.69 | 58.46 |
| Ligand |  | 62.28 |
| R.m.s. deviations |  |  |
| Bond lengths (Å) | 0.003 | 0.004 |
| Bond angles (°) | 0.551 | 0.552 |
| Validation |  |  |
| MolProbity score | 1.71 | 1.58 |
| Clashscore | 8.58 | 4.91 |
| Poor rotamers (%) | 0.11 | 0.00 |
| Ramachandran plot |  |  |
| Favored (%) | 96.31 | 95.31 |
| Allowed (%) | 3.69 | 4.69 |
| Disallowed (%) | 0.00 | 0.00 |
| **Model coverage** |  |  |
| Chain A | V31-K50; Y69-R74;<br>T76-F102; P112-V335;<br>G337-L895;<br>M906-E919 | A4-T896; M902-Q932 |
| Chain B | E77-A191 | M67-N192 |
| Chain C | K2-L71 | S1-E73 |
| Chain D | T84-I132 | T84-L122; K127-A191 |

## References and Notes

1. J. F. Chan et al., A familial cluster of pneumonia associated with the 2019 novel coronavirus indicating person-to-person transmission: a study of a family cluster. Lancet (London, England) 395, 514–523 (2020).

2. F. Wu et al., A new coronavirus associated with human respiratory disease in China. Nature, (2020).

3. N. Chen et al., Epidemiological and clinical characteristics of 99 cases of 2019 novel coronavirus pneumonia in Wuhan, China: a descriptive study. Lancet (London, England) 395, 507–513 (2020).

4. P. Zhou et al., A pneumonia outbreak associated with a new coronavirus of probable bat origin. Nature, (2020).

5. J. Ziebuhr, The coronavirus replicase. Curr Top Microbiol Immunol 287, 57–94 (2005).

6. R. N. Kirchdoerfer, A. B. Ward, Structure of the SARS-CoV nsp12 polymerase bound to nsp7 and nsp8 co-factors. Nat Commun 10, 2342 (2019).

7. M. Wang et al., Remdesivir and chloroquine effectively inhibit the recently emerged novel coronavirus (2019-nCoV) in vitro. Cell Res, (2020).

8. M. L. Holshue et al., First Case of 2019 Novel Coronavirus in the United States. N Engl J Med, (2020).

9. K. C. Lehmann et al., Discovery of an essential nucleotidylating activity associated with a newly delineated conserved domain in the RNA polymerase-containing protein of all nidoviruses. Nucleic acids research 43, 8416–8434 (2015).

10. Y. Zhai et al., Insights into SARS-CoV transcription and replication from the structure of the nsp7-nsp8 hexadecamer. Nature structural & molecular biology 12, 980–986 (2005).

11. S. M. McDonald, RNA synthetic mechanisms employed by diverse families of RNA viruses. Wiley Interdiscip Rev RNA 4, 351–367 (2013).

12. T. C. Appleby et al., Viral replication. Structural basis for RNA replication by the hepatitis C virus polymerase. Science 347, 771–775 (2015).

13. P. Gong, O. B. Peersen, Structural basis for active site closure by the poliovirus RNA-dependent RNA polymerase. Proceedings of the National Academy of Sciences of the United States of America 107, 22505–22510 (2010).

14. P. V. Afonine et al., New tools for the analysis and validation of cryo-EM maps and atomic models. Acta Crystallogr D Struct Biol 74, 814–840 (2018).

15. T. K. Warren et al., Therapeutic efficacy of the small molecule GS-5734 against Ebola virus in rhesus monkeys. Nature 531, 381–385 (2016).

16. E. J. Gane et al., Nucleotide polymerase inhibitor sofosbuvir plus ribavirin for hepatitis C. N Engl J Med 368, 34–44 (2013).

17. E. S. Svarovskaia et al., Infrequent development of resistance in genotype 1-6 hepatitis C virus-infected subjects treated with sofosbuvir in phase 2 and 3 clinical trials. Clinical infectious diseases : an official publication of the Infectious Diseases Society of America 59, 1666–1674 (2014).

18. Z. Jin et al., Structure of Mpro from COVID-19 virus and discovery of its inhibitors. bioRxiv, 2020.2002.2026.964882 (2020).

19. H. Yang et al., Design of wide-spectrum inhibitors targeting coronavirus main proteases. PLoS Biol 3, e324 (2005).

20. H. Yang et al., The crystal structures of severe acute respiratory syndrome virus main protease and its complex with an inhibitor. Proceedings of the National Academy of Sciences of the United States of America 100, 13190–13195 (2003).

21. D. N. Mastronarde, Automated electron microscope tomography using robust prediction of specimen movements. J. Struct. Biol. 152, 36–51 (2005).

22. S. Q. Zheng et al., MotionCor2: anisotropic correction of beam-induced motion for improved cryo-electron microscopy. Nat. Methods. 14, 331 (2017).

23. A. Punjani, J. L. Rubinstein, D. J. Fleet, M. A. Brubaker, cryoSPARC: algorithms for rapid unsupervised cryo-EM structure determination. Nat. Methods. 14, 290 (2017).

24. R. N. Kirchdoerfer, A. B. Ward, Structure of the SARS-CoV nsp12 polymerase bound to nsp7 and nsp8 co-factors. Nat Commun 10, 2342 (2019).

25. S. Li et al., New nsp8 isoform suggests mechanism for tuning viral RNA synthesis. Protein Cell 1, 198–204 (2010).

26. E. F. Pettersen et al., UCSF Chimera--a visualization system for exploratory research and analysis. J Comput Chem 25, 1605–1612 (2004).

27. P. Emsley, B. Lohkamp, W. G. Scott, K. Cowtan, Features and development of Coot. Acta Crystallogr D Biol Crystallogr 66, 486–501 (2010).

28. P. V. Afonine et al., Towards automated crystallographic structure refinement with phenix.refine. Acta Crystallogr D Biol Crystallogr 68, 352–367 (2012).

